# Ankyrins are essential at the photoreceptor synapse in the mouse outer retina

**DOI:** 10.1101/2025.02.11.637690

**Authors:** Ross M Perez, Jay Campbell, Martina Cavallini, Collin Todora, Daniela Becerril, Debalina Goswami-Sewell, Rajashree Venkatraman, Cesiah C Gomez, Caitlin Bagnetto, Audrey Lee, Marlon F Mattos, Yi-Rong Peng, Mrinalini Hoon, Elizabeth Zuniga-Sanchez

## Abstract

Retinal circuit assembly relies on the precise timing and positioning of key molecules between neuronal partners to mediate proper synapse formation. In the outer retina, horizontal cells are important interneurons that make the first contacts to photoreceptors and begin to segregate visual information into two distinct pathways by selectively forming synapses to the different types of photoreceptors. Dendrites of horizontal cells synapse exclusively to cone photoreceptors whereas the axon terminal synapses to rod photoreceptors. Failure to properly form these early connections disrupts the downstream connectivity of other postsynaptic neurons and leads to abnormal visual function. Although these early events are critical for proper synapse development, little is known about the molecular mechanisms that establish horizontal cell to photoreceptor connectivity during development. In the present study, we performed single-cell RNA sequencing and uncovered new molecules that are highly expressed in horizontal cells. These include different members of the cytoskeletal scaffolding family of Ankyrins that are known to form specialized regions within neurons by recruiting different molecules to the membrane and linking them to the cytoskeleton. Specifically, we found Ankyrin-B to be highly expressed in horizontal cells at early time points and Ankyrin-G to be expressed at later stages. Loss of both Ankyrin-B and Ankyrin-G leads to synaptic defects between horizontal cells and photoreceptors and disrupts *in vivo* retinal responses. In summary, our findings uncovered a new role for Ankyrins in mediating synaptic connectivity between horizontal cells and photoreceptors required for normal visual function.

**SIGNIFICANCE STATEMENT:** In the mammalian retina, the first synapse between photoreceptors and their downstream targets begins to separate visual information into two distinct pathways. During retinal development, photoreceptors first make contacts to horizontal cells in a temporal- and spatial-specific manner. Although this initial contact is critical for synaptogenesis, little is known about the key molecules responsible for selective wiring of horizontal cells to photoreceptors. In this study, we performed single-cell RNA sequencing and identified the family of cytoskeletal scaffolding proteins Ankyrins to be differentially expressed in horizontal cells. Loss of Ankyrins impairs synaptic connectivity between horizontal cells and their photoreceptor partners leading to abnormal visual responses. Taken together, our work uncovered a new function of Ankyrins at photoreceptor synapses.

## INTRODUCTION

The first synapse in the mammalian outer retina begins to process visual information into two distinct pathways: cone- and rod-driven vision. This is largely due to the distinct connectivity patterns between cone and rod photoreceptors to downstream synaptic targets such as horizontal cells and bipolar cells. Cone photoreceptors synapse selectively to the dendrites of horizontal cells and cone bipolars to mediate daylight vision and color perception (1). Rod photoreceptors synapse to the axon terminal of horizontal cells and the dendrites of rod bipolars to facilitate night vision or low-light illumination (1). See **Figure 1A**. During retinal development, photoreceptors first make contacts to horizontal cells before making connections to bipolar cells. Horizontal cells first extend neuronal processes around postnatal day 3 (P3) and these are in close contact with the terminals of cone photoreceptors (2, 3). See **Figure 1B**. By P5, horizontal cells extend their axon (4), and from P6-8 the axon terminal begins to make contacts to rod photoreceptors (5–7) as shown in **Figure 1B**. From P9-13, dendrites of bipolars begin to innervate the terminals of photoreceptors and form synapses with their appropriate targets (3) (**Figure 1B**). By P21, synapse formation in the outer retina is largely complete.

**Figure 1:**
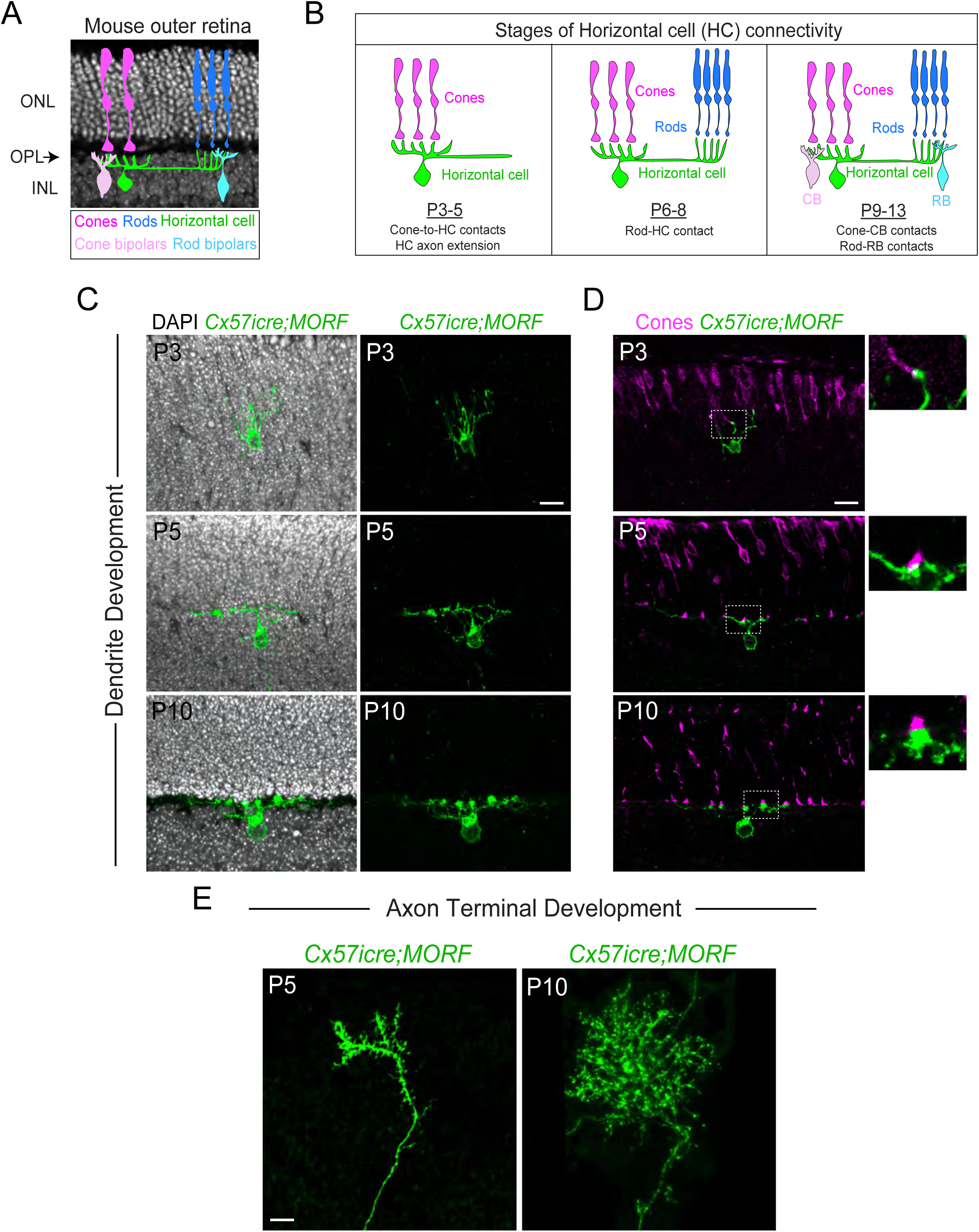
Horizontal cells undergo morphological changes during key developmental timepoints. **(A-E).** Schematic drawing of the adult mouse retina (A-B). The retina consists of three nuclear layers (ONL, INL, GCL) and two synaptic layers (OPL, IPL). Cone photoreceptors (dark pink) synapse with the dendrites of horizontal cells (green) and the dendrites of cone bipolars (light green) (A). Rod photoreceptors (blue) synapse with the axons of horizontal cells (light pink) and the dendrites of rod bipolar cells (cyan) (A). Horizontal cells first contact cone photoreceptors via their dendrites followed by axon elongation from P3-5 (B). By P6-8, the axon terminal of horizonal cells begins to contact rod photoreceptors. This is followed by innervation of the bipolar dendrites around P9-13 (B). Horizontal cells (anti-V5, green) are individually labeled using the *Tg(Cx57icre;MORF)* mouse line at distinct developmental timepoints: P3, P5, and P10 (C). DAPI was used as a nuclear stain in (C). Cone photoreceptors are labeled with anti-Sopsin at P3 and with anti-CAR at P5 and P10 in (D). Dendrite development of horizontal cells begins with contacts to cone photoreceptors (magenta) starting at P3 (D). These contacts mature from P5 to P10 (D). Insets are zoomed images of dotted boxed region in (D). The axon terminal of horizontal cells matures or extends processes from P5 to P10 (E). Scale bar = 10μm. Abbreviations: ONL=outer nuclear layer, INL=inner nuclear layer, GCL=ganglion cell layer, OPL=outer plexiform layer, IPL= inner plexiform layer.

This initial contact between photoreceptors and horizontal cells has been shown to be required for synapse formation and maintenance in the mouse outer retina. Studies of *Lim1* conditional knockouts animals where horizontal cells fail to migrate (8), do not form contacts with photoreceptors and leads to connectivity defects between rods and rod bipolars (9). These findings show horizontal cells are required for the dendrites of rod bipolars to properly innervate and form synapses with rod photoreceptors. Similarly, ablating horizontal cells at later time points after migration also disrupts connectivity between cone photoreceptors and cone bipolars which ultimately leads to photoreceptor degeneration (10). All together, these data demonstrate horizontal cells are not only required at early stages to form proper synaptic connections but also at later stages to maintain photoreceptor connectivity and survival.

Although horizontal cells play a critical role in circuit assembly, little is known about the molecular mechanisms that mediate selective wiring of horizontal cells to their respective photoreceptor target. Previous studies have shown cell adhesion molecules play a vital role in synapse formation by serving as recognition molecules to mediate interactions between pre- and postsynaptic neurons (11). Over the last decade, several cell adhesion molecules have been identified to mediate photoreceptor connectivity in the outer retina (reviewed in (1, 12)). The expression of these cell adhesion molecules is often restricted to particular cell types or specific subcellular domains to facilitate selective interactions between synaptic partners. For instance, the cell adhesion molecule Elfn1 is highly expressed in rod photoreceptors and mediates connectivity to rod bipolars (13), whereas Elfn2 is found in cone photoreceptors and mediates connectivity in the cone pathway (14). Similar restricted expression of cell adhesion molecules has also been observed within horizontal cells. The cell adhesion molecule, Ngl-2 is localized to the axon terminal of horizontal cells, and loss of Ngl-2 leads to connectivity defects between rod photoreceptors and horizontal cells (15). However, to date, Ngl-2 has been the only molecule implicated in selective wiring of horizontal cells to their appropriate photoreceptor partner.

In the present study, we set out to identify new molecules responsible for mediating synaptic connectivity between horizontal cells and photoreceptors. First, we carried out a developmental analysis using a stochastic labeling approach to identify the different subcellular compartments of horizontal cells such as the dendrites and axon terminal, and assess when they make contacts to the different types of photoreceptors. Second, we performed single-cell RNA sequencing (scRNA-seq) of horizontal cells at early developmental time points when they are actively making contacts to their respective photoreceptor partner. From our sequencing data, we found different members belonging to the cytoskeletal scaffolding Ankyrin family to be temporally expressed within horizontal cells. We found Ankyrin-B to be highly expressed at early developmental stages in horizontal cells whereas Ankyrin-G is expressed at later stages. Moreover, loss of both Ankyrin-B and Ankyrin-G disrupts synaptic connectivity between horizontal cells and photoreceptors, leading to impaired *in vivo* retinal responses in both the rod and cone pathways. In summary, our findings demonstrate a new role for Ankyrin-B and Ankyrin-G in horizontal cell connectivity required for normal visual function.

## MATERIALS AND METHODS

All mouse experiments were approved by the Institutional Animal Care and Use Committee at BCM. Eyes were collected at various developmental stages and processed for antibody staining or *in situ* hybridization as previously described (16). See **Table S4** for list of reagents. Confocal images were acquired using a Zeiss LSM 800 microscope. Technical replicates from different retinal sections are shown as small circles and biological replicates from different animals as big circles in all graphs. Quantification of presynaptic and postsynaptic marker expression was performed using the Imaris software as previously described (16, 17). Since CtBP2 protein does not form a puncta-like structure, we measured the maximum fluorescence intensity within the OPL using ImageJ. Values were normalized to account for surface area variability of the OPL between animals. Statistical tests were performed using the average of the three retinal sections per animal for each group. Eye cups were prepared for serial block-face scanning electron microscopy (SBFSEM) from P30 animals as described previously (18). Photoreceptor terminals and associated ribbon structures were identified according to known ultrastructural features as done previously (19, 20). Scotopic and photopic ERGs were recorded as described previously (16, 17). Photopic ERG responses were elicited by using a paired-flash protocol as described (16, 21). All statistical analysis were performed using GraphPad Prism version 9 with more details on the statistical tests and p-values given in the text and figure legends.

## RESULTS

### Developmental analysis of horizontal cell specificity to photoreceptors

To investigate the developmental mechanisms underlying horizontal cell (HC) connectivity to the different photoreceptors, we sparsely labeled HCs by crossing *Tg(Cx57icre)* mice (22) with the *Tg(MORF)* reporter line (4). This strategy allowed us to visualize individual horizontal cells, including their dendrites and axon terminals, across key developmental time points (**Figure 1C-E**). Consistent with prior studies (2, 3), we found dendrites of horizontal cells are initially vertical at P3 but then transition to lateral processes from P5 to P10, as shown in **Figure 1C**. At P3, dendrites of horizontal cells begin contacting cone photoreceptors, and these interactions continue to P10 (**Figure 1D**). Notably, these changes in HC morphology are accompanied by distinct changes in photoreceptor terminal morphology, including a flattening and enlargement at the base of the cone terminal by P10, suggesting the onset of mature synaptic contacts. Our whole-mount retinal images of *Tg(Cx57icre;MORF)* animals revealed the axon terminal of HCs begins to extend processes starting at P5 and continues to P10 (**Figure 1E**), which is consistent with previous work showing horizontal cells make contacts to rod photoreceptors during these developmental time points. The temporally coordinated morphological changes of horizontal cells led us to hypothesize that dynamic transcriptional changes accompany their structural remodeling and connectivity. To explore this further, we performed single-cell RNA sequencing (scRNA-seq) at defined developmental stages to capture the molecular programs driving HC connectivity to photoreceptors.

### Transcriptomic analysis of horizontal cells during photoreceptor connectivity

To better understand the molecular mechanisms underlying HC connectivity to photoreceptors, we performed single-cell RNA sequencing (scRNA-seq) at postnatal day 6 (P6), P8, and P11. These timepoints correspond to early (P6), mid (P8), and late (P11) stages of HC-photoreceptor connectivity. We purified horizontal cells via fluorescence-activated cell sorting (FACS) using *Tg(Cx57icre;Ai14)* animals. Next, we performed scRNA-seq using the 10x Genomics platform (23). See **Figure 2A**. After quality control and removal of non-horizontal cells based on the absence of HC marker expression (i.e. *Lhx1*, *Onecut1*, *Slc32a*, *Pax6*), we obtained a total of 11,034 high-quality horizontal cells across all developmental time points (**Figure 2B**, **Supplemental Figure 1**). Although all cells were grouped into a single cluster using unsupervised-based clustering methods, cells from each age were slightly segregated in UMAP space (see **Figure 2C**). We then performed differential expression (DE) analysis between the different ages: P6 vs P8, P8 vs P11, and P6 vs P11. See **Table S1-S3**. Significant DE genes were identified based on an adjusted p-value threshold of 0.05 and a log2 fold change ≥ 1.5. This resulted in 408 genes at P6, 114 genes at P8, and 61 genes at P11 (**Figure 2D**). Gene ontology (GO) enrichment analysis was then performed on significant DE genes, which showed differences in the most enriched terms across the different time points (**Figure 2D**). The GO term “synapse assembly” was significant at P6 and “positive regulation of neuron projection development” at P8, whereas “negative regulation of locomotion” was found at P11. These GO terms are consistent with our developmental analysis showing changes in HC morphology at early stages (P3-10).

**Figure 2:**
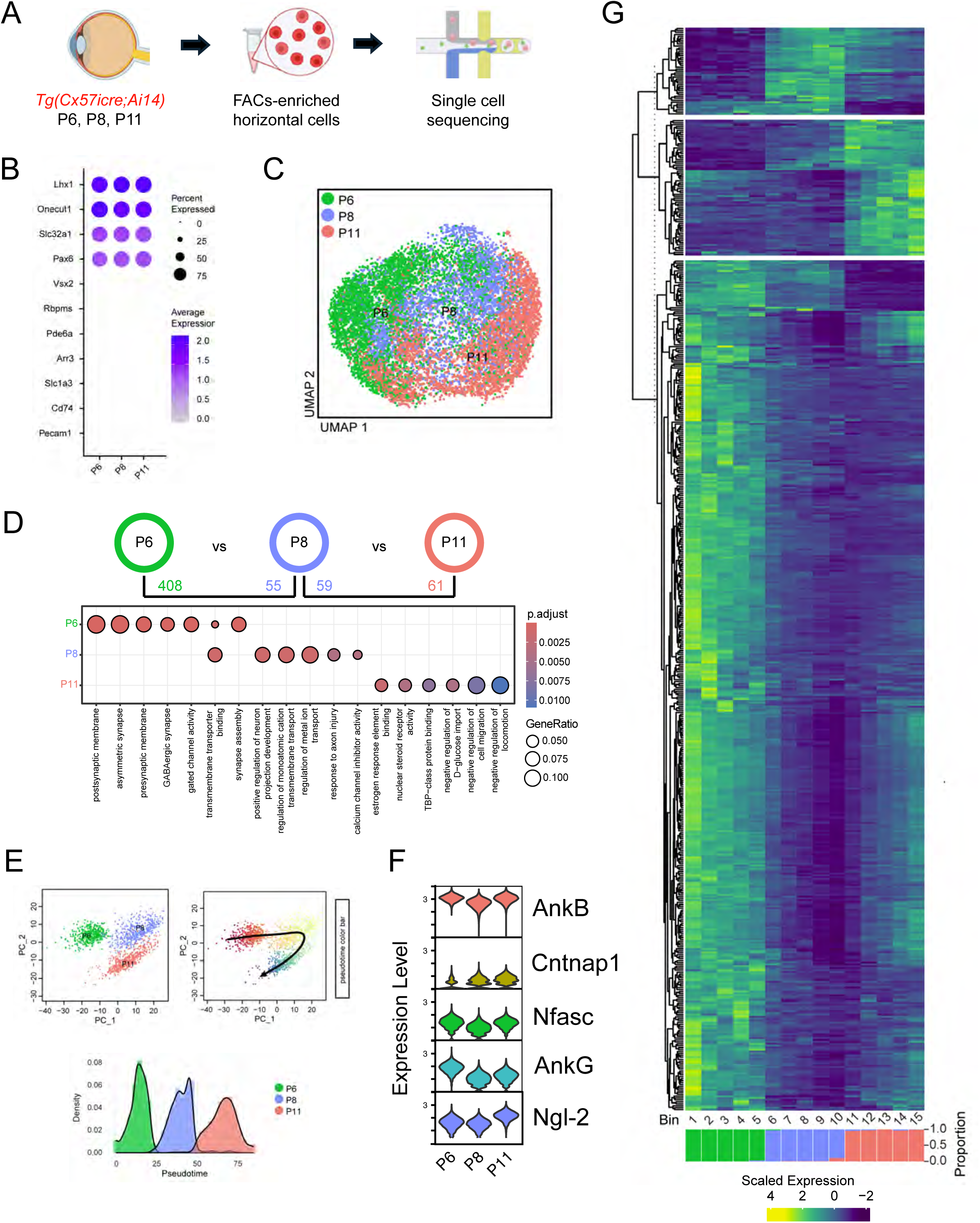
Transcriptomic data analysis and developmental trajectories of horizontal cells across different time points. **(A-G).** Schematic drawing created with BioRender.com illustrates the process of acquiring single-cell RNA sequencing data from horizontal cells (A). Dot expression profile plot of known retinal cell markers: *Lhx1*, *Onecut1*, *Slc32a1*, *Pax6* for horizontal cells, *Vsx2* for bipolars, *Rbpms* for ganglion cells, *Pde6a* for rods, *Arr3* for cones, *Slc1a3* for Muller glia, *Cd74* for microglia, and *Pecam1* for endothelial cells (B). A Uniform Manifold Approximation and Projection (UMAP) visualization of 11,034 horizontal cells with cells colored by age is shown in (C). Differential expression (DE) and gene ontology (GO) enrichment analyses were performed between adjacent ages (D). The number of significant DE genes between ages was determined using the following criteria: p.adj < 0.05 and avg_log2FC > 1.5. The number of DE genes at each time point along with the associated GO terms are shown in (D). Principal component analysis (PCA) was performed on 500 randomly sampled cells from each age group (1,500 cells total). The top left panel shows cells colored by age, while the top right panel shows the same cells colored by pseudotime, as determined by Slingshot trajectory analysis. The bottom panel shows the distribution of cells from each age group across pseudotime (E). Expression of candidate genes across experimental timepoints in violin plots (F). A heatmap displays the scaled expression of DE genes identified (G).

We then performed a pseudotime analysis to characterize the developmental trajectory of HCs at these different ages (**Figure 2E**). This involved performing a principal component analysis (PCA) of 500 randomly sampled HCs from each time point (1,500 cells total) (**Figure 2E**). To visualize pseudotime progression, cells were plotted in PCA space and colored by their pseudotime values. The results revealed distinct temporal expression patterns corresponding to different phases of HC development. To analyze gene expression changes along the developmental trajectory, we binned cells based on their pseudotime values. The scaled expression values were visualized as a heatmap with genes clustered into the three distinct age groups (**Figure 2G**). We found the cell adhesion molecule, *Ngl-2*, in our dataset, which is known to mediate HC-rod connectivity along with other cell adhesion molecules such as *Cntnap1* and *Nfasc* (**Figure 2F**). Moreover, we also identified different members of the cytoskeletal scaffolding Ankyrin family known to scaffold a variety of membrane proteins including cell adhesion molecules, receptors, and ion channels to mediate synapse formation in other regions of the central nervous system (reviewed in (24)). Our sequencing data uncovered promising candidate genes that may be responsible for the selective wiring of horizontal cells to their respective photoreceptor target.

### Distinct spatial and temporal expression of HC-specific candidate genes

We validated our HC sequencing data by analyzing the expression of our candidate genes across the three key developmental stages: P3, P5, and P10. We found Contactin-associated protein-like 1 (Cntnap1) to be localized to the somatodendritic region of horizontal cells starting at P3 and continues to P10 (**Figure 3A**). These findings were consistent with published data showing Cntnap1 is expressed in horizontal cells (25). However, Neurofascin (Nfasc) which is known to mediate rod connectivity (17), is expressed starting at P5 and appears to be confined to the axonal region of horizontal cells (**Figure 3B**). Previous research has shown localized expression of cell adhesion molecules mediates selective wiring of horizontal cells to photoreceptors (15). Moreover, Ankyrins are adaptor proteins that scaffold a variety of membrane proteins including cell adhesion molecules to different cellular compartments to mediate synapse formation in other regions of the central nervous system (24). Thus, we set out to investigate the role of Ankyrins in mediating horizontal cell connectivity during retinal development.

**Figure 3:**
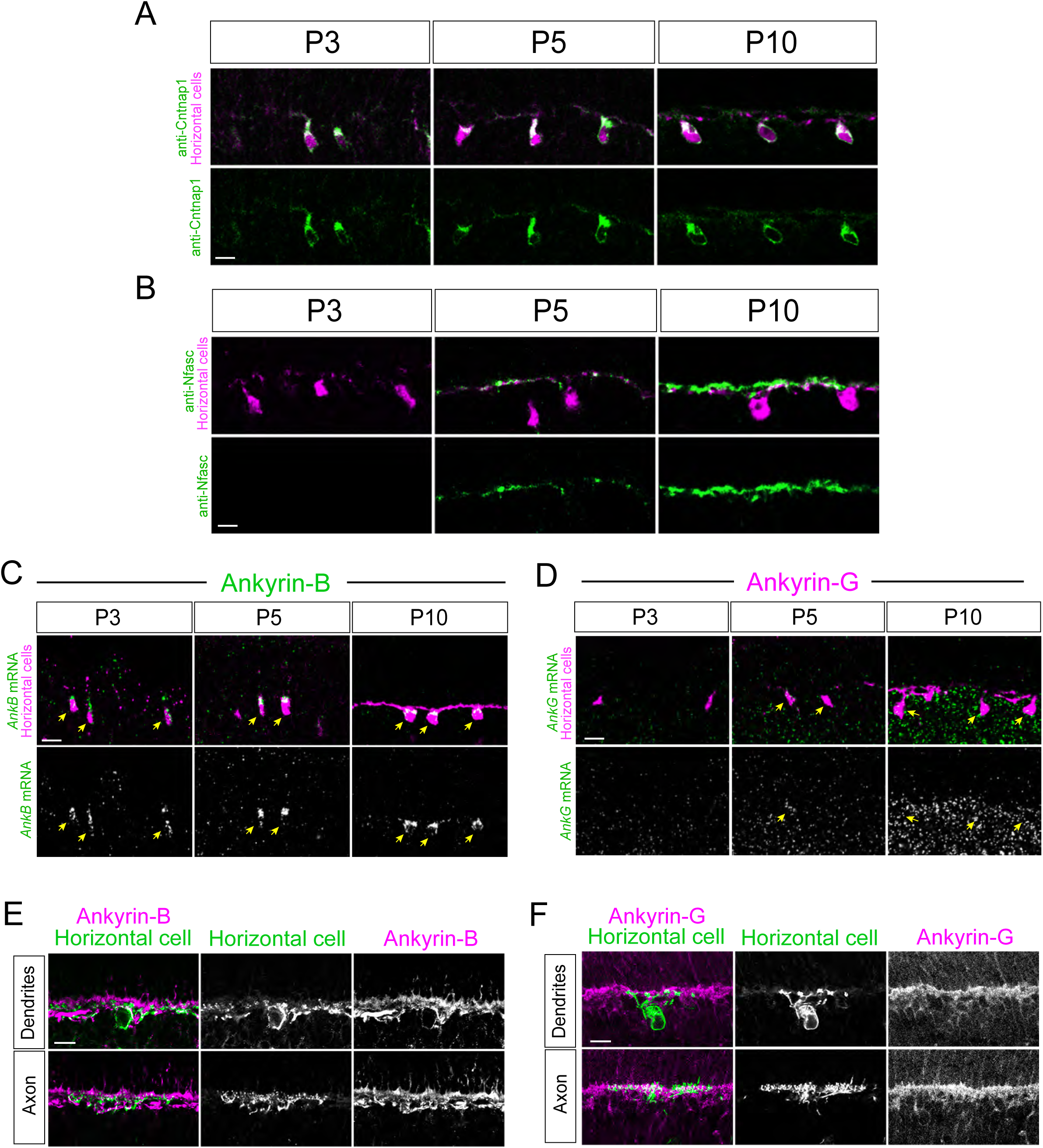
Validation of candidate genes at key developmental timepoints. **(A-F).** Protein expression of the cell adhesion molecules Cntnap 1 and Nfasc are shown in green and horizontal cells are labeled with anti-calbindin (magenta) at P3, P5, P10 in (A,B). *AnkB* and *AnkG* mRNA expression (green) was detected via *in situ* hybridization and horizontal cells were labeled with anti-calbindin (magenta) at P3, P5, P10 in (C,D). Yellow arrows denote *AnkB* or *AnkG* mRNA expression within horizontal cells in (C,D). Sparse labeling of horizontal cells at P15 using *Tg(Ptf1a-cre;MORF)* animals revealed Ankyrin-B (magenta) is found in the dendrites and axon terminal of horizontal cells (green) in (E). Similarly, Ankyrin-G (magenta) is also highly expressed in the OPL and co-localizes with the dendrites and axon terminal of horizontal cells shown in green (F). Scale bar = 10μm.

First, we examined Ankyrin-B (AnkB) and Ankyrin-G (AnkG) expression in horizontal cells at early developmental stages. By performing *in situ* hybridization followed by antibody staining, we confirmed *AnkB* mRNA is expressed in horizontal cells from P3 to P10 as shown in **Figure 3C**. However, *AnkG* mRNA is not detected in horizontal cells until P5-10 and continues to be expressed at lower levels compared to *AnkB* (**Figure 3D**). At P15, Ankyrin-B and Ankyrin-G protein is localized to the synaptic layer of the outer retina or OPL and co-localizes with the dendrites and axon terminal of horizontal cells (**Figure 3E-F**). By P30, Ankyrins continue to be highly expressed in the OPL and showed extensive overlap with processes from horizontal cells and their respective synaptic partners (**Supplemental Figure 2A-D**). These findings indicate Ankyrins are not only expressed during key stages of horizontal cell connectivity but are also localized to the synaptic layer where horizontal cells form synapses to their respective photoreceptor partners.

### Loss of Ankyrins leads to horizontal cell sprouts and OPL defects

We then performed functional studies by crossing published floxed alleles of Ankyrin-B (*AnkB^flox/flox^*) (26) and Ankyrin-G (*AnkG^flox/flox^*) (27) to the early retinal progenitor cell driver, *Chx10-cre* (28), to conditionally delete Ankyrins, as germline knockouts are known to be embryonically lethal (29). This resulted in *Chx10-cre;AnkB^flox/flox^* and *Chx10-cre;AnkG^flox/flox^*transgenic animals which we refer to as AnkB CKO and AnkG CKO, respectively. We confirmed loss of Ankyrin protein expression in CKO animals by performing antibody staining as shown in **Supplemental Figure 2E**. Next, we examined AnkB CKO and AnkG CKO for morphological and synaptic defects using available antibodies. By P30, processes from horizontal cells seen with calbindin staining are normally confined to the OPL in wild-type retinas as shown in controls (**Figure 4A**), and misprojections or sprouts from horizontal cells into the ONL are correlated with synaptic defects in diseased and aged mouse retinas (15, 16, 30, 31). We found single knockouts of Ankyrin-B or Ankyrin-G led to minimal horizontal cell sprouts compared to controls (**Figure 4A**). Moreover, expression of a known presynaptic marker, anti-CtBP2 localized to both rod and cone terminals, appears unaffected in AnkB CKO and AnkG CKO compared to controls (**Figure 4A**). As Ankyrin-B and Ankyrin-G are both expressed in horizontal cells based on our sequencing data (**Figure 2F**), and they are known to be functionally redundant with one another (32), we generated *Chx10-cre;AnkB^flox/flox^;AnkG^flox/flox^* double knockout animals which we refer to as AnkB; AnkG DKO. We confirmed Ankyrin-B is still present in AnkG CKO and Ankyrin-G in AnkB CKO; however, only both are significantly reduced in the OPL of AnkB; AnkG DKO animals (**Supplemental Figure 2E**). We found disruption of both Ankyrin-B and Ankyrin-G results in numerous misprojections or sprouts from horizontal cells extending beyond the OPL into the nuclear layer or ONL as depicted by arrows in **Figure 4A**. Consistent with misprojections of horizontal cells associated with synaptic defects, we observed a significant decrease in the presynaptic marker, CtBP2 within the OPL of AnkB; AnkG DKO compared to controls (**Figure 4A**). These observations were quantified by measuring the total volume of calbindin staining in the ONL and CtBP2 immunofluorescence signal in the OPL (**Figure 4D**). Controls and single knockout animals showed minimal calbindin staining in the ONL; however, AnkB; AnkG DKO displayed significant sprouts from horizontal cells with reduced CtBP2 expression in the OPL (**Figure 4A,D**). Next, we further examined synaptic changes due to loss of Ankyrins. Similar to our CtBP2 findings in **Figure 4A**, we found reduced immunofluorescence signal of the presynaptic marker, Bassoon (Bsn) in the OPL of AnkB; AnkG DKO animals compared to controls (**Figure 4B**). Bsn is normally expressed in both rod and cone terminals and forms puncta-like structure that is about 0.6μm in diameter (16, 17). Using the Imaris confocal software, we quantified the total number of Bsn puncta in the OPL and found an 18% reduction in the OPL of AnkB; AnkG DKO compared to controls (**Figure 4D**). In addition, we also examined changes of postsynaptic receptors such as the metabotropic glutamate receptor 6 (mGluR6) that is localized to the dendrites of ON bipolars (33, 34), and found a 28% reduction of mGluR6 in AnkB; AnkG DKO compared to controls (**Figure 4C**). These results show reduced presynaptic and postsynaptic immunofluorescence signal in the OPL due to loss of Ankyrins.

**Figure 4:**
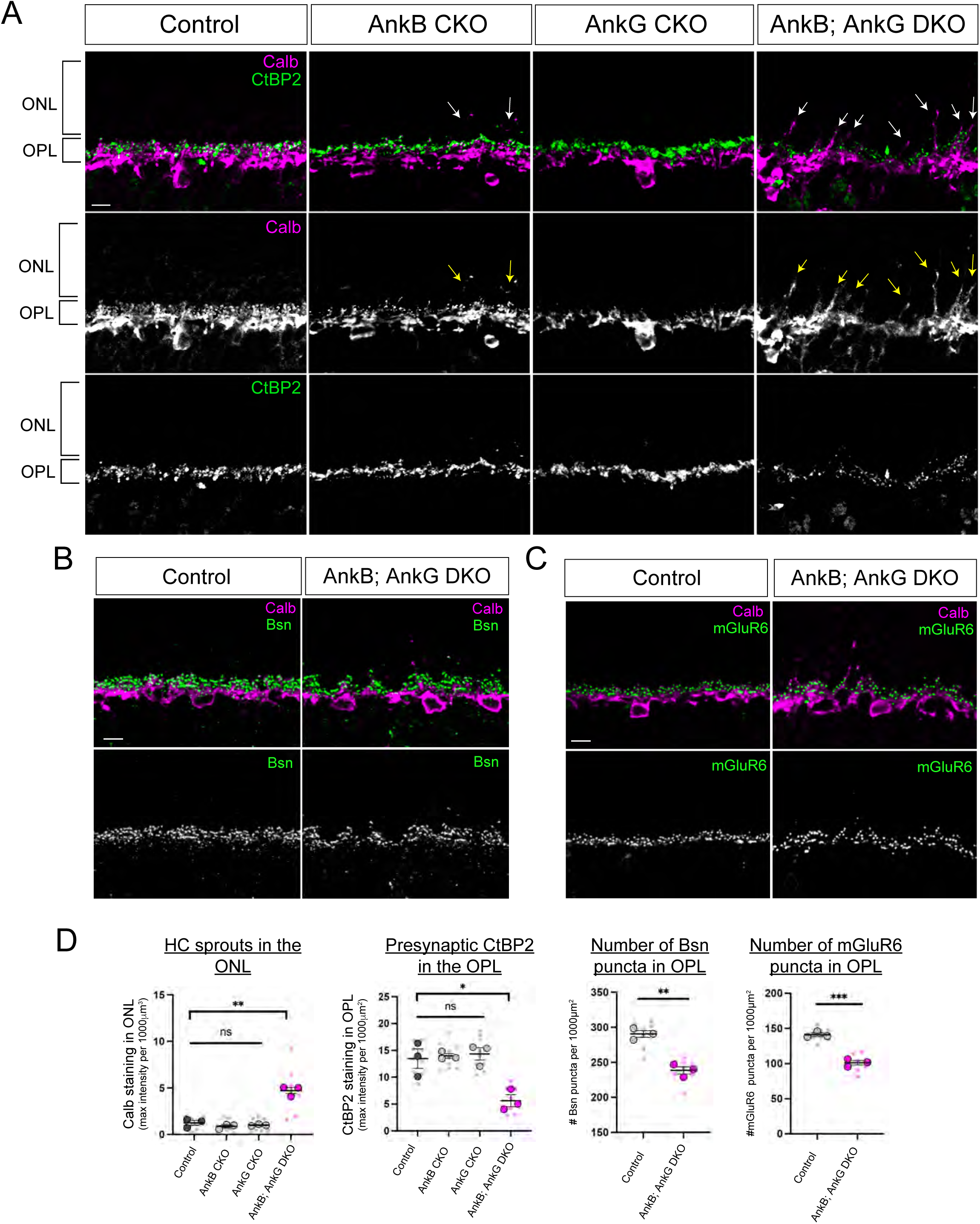
Loss of Ankyrin-B and Ankyrin-G results in horizontal cell sprouts and OPL defects. **(A-C).** Processes from horizontal cells labeled with anti-calbindin (Calb, magenta) are normally confined to the OPL by P30 as shown in controls (A). Single conditional knockouts of Ankryin-B (AnkB CKO) or Ankyrin-G (AnkG CKO) leads to minimal misprojections or sprouts of horizontal cells (A). However, disruption of both Ankyrin-B and Ankyrin-G or AnkB; AnkG DKO results in multiple horizontal cell sprouts in the ONL as depicted by arrows in (A). Expression of the known presynaptic marker, CtBP2 (green) showed a significant reduction in the OPL in AnkB; AnkG DKO but not in AnkB CKO or AnkG CKO compared to controls (A). Protein expression of another presynaptic marker, Bassoon (Bsn, green) and the postsynaptic marker, mGluR6 (green) is also reduced in the OPL in AnkB; AnkG DKO compared to controls (B,C). Horizontal cells are labeled with anti-calbindin (Calb, magenta) in (B,C). Horizontal cell (HC) sprouts were quantified by measuring the amount of calbindin staining in the ONL in (D). CtBP2 immunofluorescence signal in the OPL was measured across all experimental groups in (D). Quantification of Bsn and mGluR6 immunofluorescence signal in the OPL was measured using the Imaris software to automatically detect puncta of about 0.6μm in size (D). The average from three different mice were used for statistical analysis. Data from each animal are represented as mean values ± SEM. Statistical significance determined by an unpaired two-tailed Student’s t test. *p < 0.05, **p < 0.01, ***p < 0.001. Scale bar = 10μm.

To determine if these changes were due to retinal degeneration, we measured the different retinal layers in both control and AnkB; AnkG DKO animals at P30 (**Supplemental Figure 3A-B**). We found a slight reduction in the ONL but not the INL or GCL of AnkB; AnkG DKO animals compared to controls. Next, we assessed the role of Ankyrins on photoreceptors by using antibodies that label the outer segments and axon terminals of rod and cone photoreceptors. We used anti-CNGA1 and PNA staining to label the outer segments of rods and cones, respectively (**Supplemental Figure 4A-D**). We found no significant difference between the rod outer segments of controls and AnkB; AnkG DKO animals (**Supplemental Figure 4B**). Similarly, there was also no significant difference in the total number of cone outer segments or their volume between controls and AnkB; AnkG DKO animals (**Supplemental Figure 4D**). We also examined the axon terminals of rod photoreceptors (anti-PSD-95) and cone photoreceptors (anti-CAR) as well as the dendrites of rod bipolars (anti-PKC) and cone bipolars (anti-Scgn). See **Supplemental Figure 4E**. Our results show there is no retraction of axon terminals from photoreceptors even in regions where there is sprouting from horizontal cells as depicted by white arrows in AnkB; AnkG DKO retinas (**Supplemental Figure 4E**). Similarly, we do not observe sprouting or mispositioning of dendrites from rod bipolars nor cone bipolars due to loss of Ankyrins (**Supplemental Figure 4E**). These findings suggest that although there is some loss of photoreceptors, Ankyrins are mainly required to maintain processes from horizontal cells into the OPL but not for other neuronal types.

### Disruption of synaptic innervation between horizontal cells and photoreceptors

To further examine the synaptic changes associated with loss of Ankyrins, we performed serial block-face scanning electron microscopy (SBFSEM) on AnkB; AnkG DKO retinas at P30. We found processes from horizontal cells (shown in yellow), which are usually confined to the cone synapse or pedicle in wild-type retinas, were extending beyond the cone terminal in AnkB; AnkG DKO, as depicted by white arrows in **Figure 5A**. These findings were similar to the horizontal cell sprouting phenotype seen with calbindin labeling in AnkB; AnkG DKO retinas in **Figure 4A**. Interestingly, dendrites of bipolar cells (cyan) appear disorganized but remain confined to the cone pedicle as depicted by white arrowheads in **Figure 5A**. These results were also consistent with our confocal data showing no sprouts from cone bipolars as seen in **Supplemental Figure 4E.** Moreover, we found the ribbons of cone pedicles in AnkB; AnkG DKO were not anchored to the base as in wild type, but rather floating as shown by asterisk in **Figure 5A**. Similar floating ribbons have been reported previously in Bassoon mutant mice, which also display synaptic defects in the outer retina (35). We quantified the percentage of ribbons associated with a horizontal cell (HC), bipolar cell (BC), glial partner, or unidentified (UI) neuronal process (**Figure 5C**). We found there is a trend towards reduced contact between cone pedicles and horizontal cells, as well as bipolar cells in the AnkB; AnkG DKO. These data were consistent with the decrease in pre- and postsynaptic immunofluorescence signal in the OPL as shown in **Figure 4B,C**.

**Figure 5:**
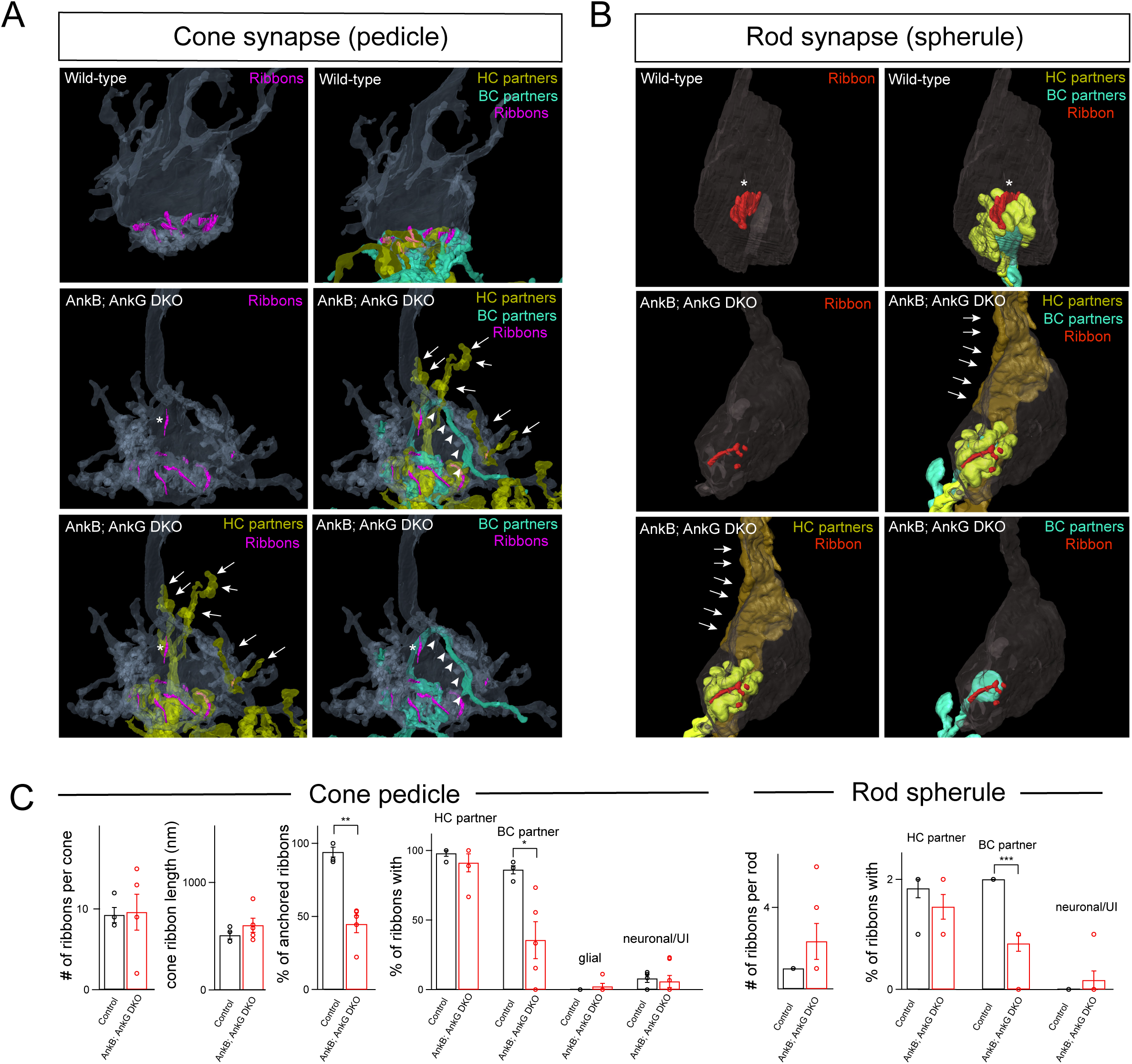
AnkB; AnkG DKO mice display aberrant synaptic innervation as seen with SBFSEM. **(A-C).** A 3D reconstruction of a wild-type and AnkB; AnkG DKO cone synapse or pedicle (grey) with included ribbons (pink), horizontal cell partners (HC, yellow), and bipolar cell partners (BC, cyan) in (A). A 3D reconstruction of a wild-type and AnkB; AnkG DKO rod synapse or spherule innervated by horizontal cell partners (HC, yellow) and bipolar cell partners (BC, cyan) in (B). Quantification of the number of ribbons per cone pedicle, the ribbon length, the % of anchored ribbon and the % of ribbons within a pedicle associated with HC partners, BC partners, glial partner and unidentified (UI) neuronal processes (C). A total of 5 cone pedicles were reconstructed from two different AnkB; AnkG DKO animals, and 4 wild-type/control cone pedicles were used for the analysis in (C). A total of 6 rod spherules were reconstructed from AnkB; AnkG DKO mice, and 6 reconstructed wild-type rod spherules were used for the analysis in (C). The % of anchored ribbons and those associated with BC partners in cone pedicles was statistically significant between wild-type and AnkB; AnkG DKO. In addition, the number of BC partners is significantly reduced for AnkB; AnkG DKO rod spherules compared to controls. Statistical significance was determined by an unpaired two-tailed Student’s t-test. *p < 0.05, **p < 0.01, ***p < 0.001.

Next, we performed a similar 3D reconstruction analysis of the rod synapse or spherule in animals with loss of Ankyrins. Rod terminals typically have one large horseshoe-shaped ribbon or synaptic site (19, 36) as shown by yellow arrow in **Figure 5B**. Our data shows that one of the rod terminals from AnkB; AnkG DKO contains multiple small, fragmented ribbons (yellow arrowhead) with one of the HC processes (shown in brown) extending beyond the synaptic layer into the ONL, as depicted by white arrows. In contrast, the other HC process (yellow) appears to properly innervate the rod spherule in **Figure 5B**. Similarly, dendrites of bipolar cells (shown in cyan) seem to be localized to the rod spherule and do not extend beyond the synaptic layer in **Figure 5B**, which is consistent with our confocal findings shown in **Supplemental Figure 4E**. Quantifications of these observations are shown in **Figure 5C**. Moreover, we also measured the total number of synaptic vesicles located at the ribbon of cone and rod synapses from wild-type and AnkB; AnkG DKO animals (**Supplemental Figure 5A-B**). Our data revealed a significant decrease in the number of vesicles of both cone and rod synapses (**Supplemental Figure 5C**). These data further support the role of Ankyrins in mediating proper synaptic innervation in the outer retina.

### Impaired rod- and cone-driven retinal responses in animals with loss of Ankyrins

We then addressed whether the synaptic defects seen in AnkB; AnkG DKO result in abnormal retinal responses. To answer this question, we performed *in vivo* full-field electroretinograms (ERG) recordings in both dark-adapted and light-adapted mice to measure rod-driven and cone-driven responses, respectively. Anesthetized mice were exposed to different flashes of light to measure rod-driven responses (scotopic ERG) or cone-driven responses (photopic ERG) as described in (16, 17, 21, 37). ERG responses are comprised of an initial a-wave response elicited by photoreceptors, followed by a b-wave response generated by postsynaptic neurons such as bipolars (37). For scotopic responses, we plotted individual ERG traces elicited from various stimulus intensities from controls (black line) and AnkB; AnkG DKO (magenta line) animals (**Figure 6A**). We quantified these observations by plotting the a-wave and b-wave amplitudes of different animals from both controls and AnkB; AnkG DKO animals as shown in **Figure 6B**. We calculated the maximum rod-driven b-wave responses or Bmax and found controls to have a Bmax value of 355 μV, whereas AnkB; AnkG DKO have a Bmax value of 217 μV. Our data confirms that the rod component of the b-wave, but not the a-wave is reduced at low light intensities in AnkB; AnkG DKO animals, and only at higher flash intensities, where there is a mixture of rod and cone responses, are a-wave responses affected due to loss of Ankyrins (**Figure 6B**). We also measured cone-driven or photopic responses in controls and AnkB; AnkG DKO animals using a paired-flash protocol (38). Individual traces of photopic ERG responses showed a significant reduction in b-wave responses (bipolars) in AnkB; AnkG DKO animals compared to controls (**Figure 6C**). Moreover, quantification of the a-wave and b-wave amplitudes of cone-driven or photopic ERG responses showed a significant reduction in AnkB; AnkG DKO animals compared to controls (**Figure 6D**). The significant reduction in b-wave responses in both the rod and cone pathways are consistent with synaptic defects between photoreceptors and their respective downstream partners.

**Figure 6:**
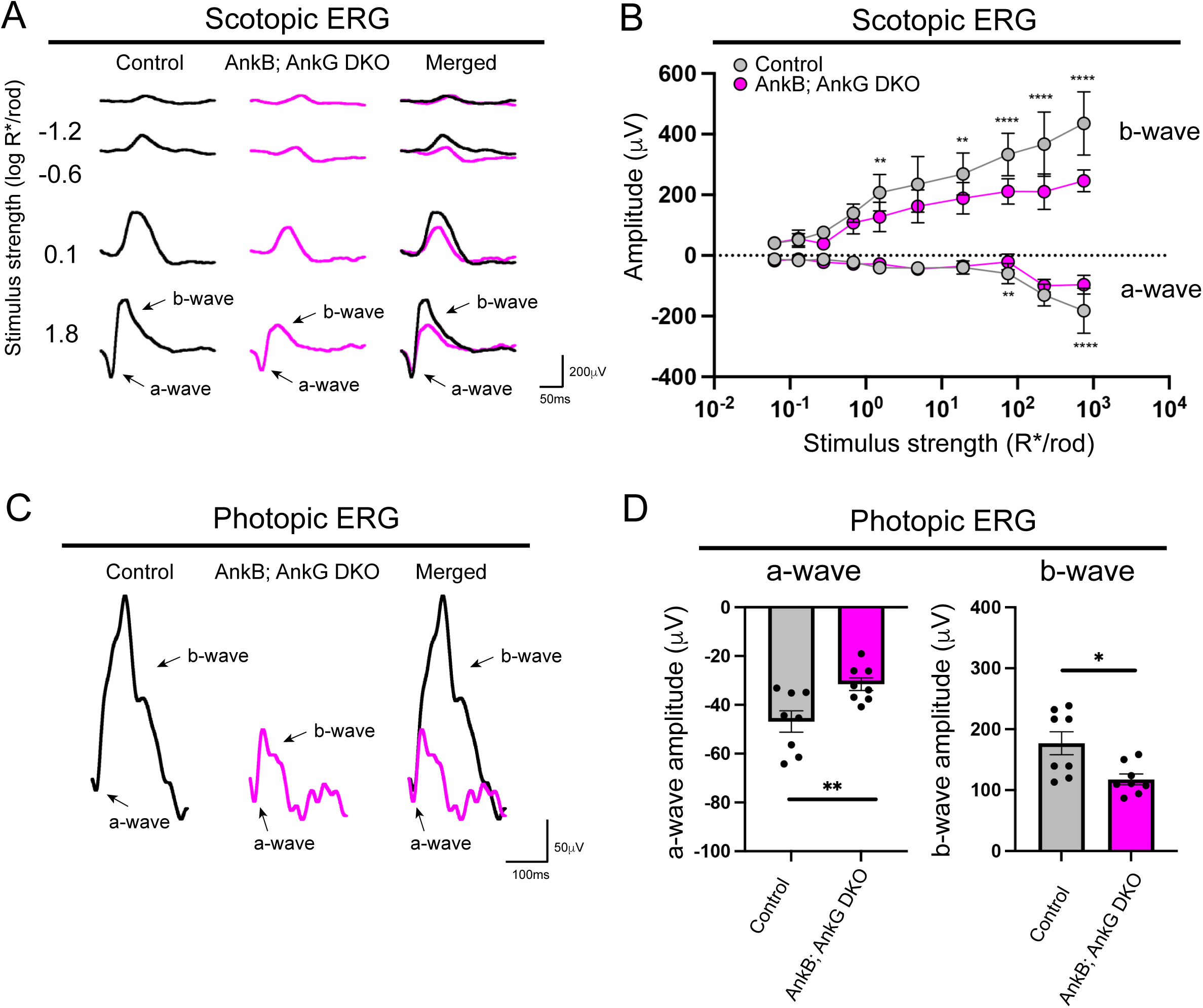
Reduced *in vivo* retinal responses in mice with loss of Ankyrin function. **(A-D).** Individual electroretinogram (ERG) traces from controls (black) and AnkB; AnkG DKO (magenta) are shown at different scotopic or rod-driven stimulus intensities (A). Scotopic b-wave amplitudes are significantly reduced in AnkB; AnkG DKO (magenta line, n=8 from four mice) compared to controls (black line, n=8 from four mice) at both low and high light stimulus in (B). However, scotopic a-wave amplitudes are only significantly reduced in AnkB; AnkG DKO at high light stimulus (B). Individual electroretinogram (ERG) traces from controls (black) and AnkB; AnkG DKO (magenta) are shown at photopic or cone-driven stimulus (C). Reduced photopic ERG responses in both a-wave and b-wave are observed in AnkB; AnkG DKO animals compared to controls using a paired flash method (n=8 from four mice per group) in (D). Data represented as mean values ± SEM. Statistical significance was determined using Student’s t-tests with Holm-Sidak correction for multiple comparisons for scotopic ERGs, and an unpaired two-tailed Student’s t-test for photopic ERGs. ns p > 0.05, *p < 0.05, **p < 0.01, ****p < 0.0001.

## DISCUSSION

Our work uncovered a new role for Ankyrins at photoreceptor synapses in the mouse outer retina. We found Ankyrin-B and Ankyrin-G are expressed in horizontal cells at early developmental stages, and protein expression is localized to the synaptic layer of the outer retina in adult mice. Moreover, loss of Ankyrin-B and Ankyrin-G leads to (1) mispositioned processes or sprouts from horizontal cells into the ONL, (2) reduced expression of pre- and postsynaptic proteins in the OPL, (3) defects in synaptic connectivity between horizontal cells and photoreceptors, and (4) impaired *in vivo* retinal responses. Taken together, our findings reveal Ankyrins are essential at the synapse in the outer retina and are required for normal visual function.

### Horizontal cells in photoreceptor circuit assembly

In the outer retina, neurons are born shortly after birth, but synaptogenesis is delayed and occurs in a stereotyped and sequential manner postnatally (3). One of the earliest events in synapse development is the formation of connections between photoreceptors and horizontal cells. These initial contacts are known to be critical for the dendrites of bipolars to properly position themselves and form synaptic connections to photoreceptors in the OPL (9). Our developmental analysis uncovered horizontal cell specificity to photoreceptors arises at early developmental stages (P3-10) prior to connectivity with bipolar cells (P9-13). After horizontal cells have made their initial contacts to photoreceptors, bipolars begin to make contacts to the rod and cone terminals. The cell adhesion molecule Ngl2 has been shown to mediate synaptic connectivity between horizontal cells and rod photoreceptors. Loss of Ngl2 results in sprouting of horizontal cells but not of dendrites from rod bipolars (15), similar to AnkB; AnkG DKO animals. However, removal or ablation of horizontal cells leads to sprouting of bipolar dendrites at later stages (approximately 3-7 months) but not at early time points (9, 10). Although there is no sprouting of bipolar cells at early developmental stages, there is a significant reduction of pre- and postsynaptic protein expression in the OPL (9, 10), suggesting impaired synapse formation due to loss of horizontal cells. These findings imply that the initial extension and positioning of bipolar dendrites to the OPL is not dependent on horizontal cells but is still required for proper synapse development. Dendrites of bipolar cells must use horizontal cells as a recognition signal and/or scaffold to then form proper synapses with photoreceptors, and sprouting of bipolar dendrites observed at later stages may be a consequence of defective retinal connectivity.

### Function of Ankyrins in the developing nervous system

In this study, we found different members of the Ankyrin family to be temporally expressed in horizontal cells. Ankyrin-B is expressed at early stages (P3), whereas Ankyrin-G is expressed at later stages from P5-10. Ankyrins are cytoskeletal scaffolding proteins that are widely expressed throughout the developing nervous system (39, 40). They are known to play a pivotal role in forming specialized structures within the nervous system, such as the axon initial segment and nodes of Ranvier, which are essential for normal neuronal function (26, 41–43). Ankyrins form and maintain these specialized structures within neurons by recruiting different cell adhesion proteins, channels, and receptors to the membrane and linking them to the cytoskeleton (44–48). In mammals, the Ankyrin family consists of three members: Ankyrin-R, Ankyrin-B, and Ankyrin-G. Previous studies have shown Ankyrin-B and Ankyrin-G are highly expressed in the mouse retina (49–51), with Ankyrin-B localized to the synaptic layers at adult stages (52). Although Ankyrin expression in the retina had been previously reported, their function at the synapse had been largely unknown.

### Role of Ankyrins in the mouse outer retina

The first synapse of the outer retina is comprised of multiple components, including scaffolding proteins, receptors, and channels (reviewed in (1)). Loss of presynaptic proteins such as Bassoon (35) or the auxiliary subunit of the calcium channel (53) disrupts synaptic connectivity in both the rod and cone pathways. This is often seen as retraction from presynaptic neurons and/or sprouts from postsynaptic neurons. However, these molecules are shared among rod and cone synapses, which raises the question of what drives their distinct connectivity. Recent advancements in RNA sequencing have begun unveiling new molecules whose expression is restricted to either rods or cones (3, 54–56). These include several cell adhesion molecules such as Elfn1 (13), Elfn2 (14), Nfasc (17), and Lrfn2 (57). Functional studies suggest restricted expression of these molecules may mediate selective wiring of photoreceptors to their respective targets, as loss of these cell adhesion molecules disrupts connectivity of one pathway but not both. Moreover, loss of some of these cell adhesion molecules does not result in sprouting phenotypes but still disrupts synapse formation between pre- and postsynaptic partners. These findings suggest photoreceptor circuit assembly relies on multiple components, with each molecule performing a distinct function. In the case of Ankyrins, they are known to bind and recruit various components to mediate neural circuit assembly including cell adhesion molecules (24). Nfasc is a cell adhesion molecule and known binding partner of Ankyrin-G (47). However, loss of Nfasc in the retina does not recapitulate the synaptic defects seen in the retinas of AnkB; AnkG DKO animals (17). Specifically, loss of Nfasc disrupts synaptic connectivity of the rod pathway but not the cone pathway (17). Since we observe connectivity defects and abnormal retinal responses in both the rod and cone pathways due to loss of Ankyrins, there must be other cell adhesion molecules that interact with Ankyrins to mediate cone connectivity. Our ultrastructure high-resolution imaging data also uncovered different ribbon phenotypes in rod and cone photoreceptors. Ribbon synapses in rods are smaller and fragmented in AnkB; AnkG DKOs compared to controls, whereas those in cones remain intact but not anchored to the cone pedicle. This could explain why we observe a reduction and not a complete absence of pre- and postsynaptic protein expression in the OPL due to loss of Ankyrins. Presynaptic proteins such as CtBP2 and Bassoon are critical components of the ribbon synapse. Loss of Bassoon results in free-floating or unanchored ribbons in rod photoreceptors (35) similar to what we observe in cone photoreceptors with loss of Ankyrins. However, although ribbons are still present in Bassoon mutants, they are not able to elicit normal retinal responses, suggesting an impairment in synaptic transmission. Similarly, we also found impairment in cone-driven responses in AnkB; AnkG DKO compared to controls, suggesting that Ankyrins are also crucial for the cone pathway. In summary, our data uncovered a new role for Ankyrins at photoreceptor synapses of the mouse outer retina. Future studies are needed to identify the different molecules recruited by Ankyrins to mediate synaptic connectivity.

## Acknowledgements

This work was supported by the National Eye Institute (F31EY035931 to RMP, R01EY033037 to EZS, R01EY035324 to YRP, T32EY007026 to MC, P30EY002520 to BCM Department of Ophthalmology, P30EY00331 to UCLA Jules Stein Eye Institute); Research to Prevent Blindness to EZS, YRP, unrestricted department grants to BCM, UCLA, UW Madison; UCLA-Caltech Medical Scientists Training Program (T32GM008042l to MC); McPherson Eye Research Institute’s Rebecca Meyer Brown/Retina Research Foundation Professorship to MH, and the UW 2020 WARF Discovery award.

## Data availability

The scRNA-Seq data generated in this study has been deposited to the Gene Expression Omnibus (GEO) under accession GSEXXXXXX. Raw confocal images have been deposited to Zenodo (XXXXXX).

