## Supplemental Appendix for "Ankyrins are essential at the photoreceptor synapse in the mouse outer retina"

**Supplemental Figure 1: Quality control analysis of scRNA-seq sequencing data (A-D).**

Violin plots showing similar distribution across ages after quality control filtering for the number of genes (A), counts (B), percent of mitochondrial RNA (C), and the percent of ribosomal RNA (D).

**Supplemental Figure 2: Ankyrin expression in the OPL of wild-type and conditional knockout animals (A-C).**

Ankyrin-B (green) and Ankyrin-G (magenta) protein expression in wild-type mice at P30 showing high levels of expression within the OPL as shown by black arrow in (A,B). DAPI staining was used to label the nuclear layers. Ankyrin-B protein (shown in green) co-localizes with rod terminals (anti-PSD-95, magenta) and cone terminals (anti-CAR, magenta), dendrites of rod bipolars (anti-PKC, magenta) and cone bipolars (anti-Scgn, magenta), and processes from horizontal cells (anti-Calb, magenta) in wild-type retinas at P30 in (C). Ankyrin-G (shown in magenta) also shows a similar expression pattern as Ankyrin-B with co-localization of pre- and postsynaptic neurons in wild-type retinas at P30 (D). Ankyrin-B protein expression (green) is reduced in the OPL of AnkB CKO and AnkB; AnkG DKO but not in AnkG CKO animals compared to controls at P30 (E). Ankyrin-G protein expression (magenta) is reduced in the OPL of AnkG CKO and AnkB; AnkG DKO but not in AnkB CKO animals compared to controls at P30 (E). Scale bar shown on images.

**Supplemental Figure 3: Loss of Ankyrins leads to a slight reduction of the ONL but not the INL or GCL (A-B).**

DAPI staining was used to label the different retinal layers in control and AnkB; AnkG DKO animals at P30 (A). There is a slight reduction of the ONL layer where the cell bodies of photoreceptors reside in AnkB; AnkG DKO compared to controls (A). However, no detectable change was observed in the INL or GCL layers (A). Measurements of the different retinal layers are shown in (B). The average of three mice per group were used for statistical analysis. Data from three different animals are represented as mean values ± SEM. Statistical significance determined by an unpaired two-tailed Student's t test. ns $p > 0.05$, *$p < 0.05$.

**Supplemental Figure 4: Loss of Ankyrins does not disrupt outer segments of photoreceptors or positioning of other neuronal subtypes (A-E).**

The outer segments of rod photoreceptors were visualized using anti-CNGA1 (magenta) from both controls and AnkB; AnkG DKO animals at P30 (A). Measurements of the CNGA1 fluorescence intensity across different animals is shown in (B). The outer segments of cone photoreceptors were detected using PNA staining (green) in controls and AnkB; AnkG DKO animals (C). The total number of cone outer segments and their respective volume is shown in (D). Data of the three different animals are represented as mean values ± SEM. Statistical significance determined by an unpaired two-tailed Student's t test. ns $p > 0.05$. DAPI staining was used to visualize the ONL in (A) and (C). Axon terminals of rod photoreceptors (anti-PSD-95) and cone photoreceptors (anti-CAR) are properly positioned to the OPL in both controls and AnkB; AnkG DKO even in regions where there is sprouting of processes from horizontal cells (anti-Calb, magenta) as depicted by arrows (E). Similarly, dendrites from rod bipolars (anti-PKC) and cone bipolars (anti-Scgn) are also confined to the OPL in both controls and AnkB; AnkG DKO animals (E). Scale bar shown on the figure.

**Supplemental Figure 5: Reduced number of synaptic vesicles in photoreceptor synapses due to loss of Ankyrins (A-C).**

Raw electron microcopy (EM) images of a single plane are shown for a cone synapse or pedicle (purple, A) and a rod synapse or spherule (yellow, B) in a wild-type and AnkB; AnkG DKO retina. The average number of synaptic vesicles found in cone pedicles and rod spherules are shown in (C). Data are represented as mean values ± SEM. Statistical significance was determined by an unpaired two-tailed Student's t test. $^*p < 0.05$, $^{**}p < 0.01$.

**Supplemental Table 1**: List of differentially expressed (DE) genes between P6 vs P8.

**Supplemental Table 2**: List of differentially expressed (DE) genes between P8 vs P11.

**Supplemental Table 3**: List of differentially expressed (DE) genes between P6 vs P11.

**Supplemental Table 4**: List of antibodies used in this study.
